# Genetic Predisposition to Neuroblastoma Results from a Regulatory Polymorphism that Promotes the Adrenergic Cell State

**DOI:** 10.1101/2023.02.28.530457

**Authors:** Nina Weichert-Leahey, Hui Shi, Ting Tao, Derek A Oldridge, Adam D Durbin, Brian J. Abraham, Mark W Zimmerman, Shizhen Zhu, Andrew C Wood, Deepak Reyon, J Keith Joung, Richard A Young, Sharon J Diskin, John M. Maris, A Thomas Look

## Abstract

Childhood neuroblastomas exhibit plasticity between an undifferentiated neural crest-like “mesenchymal” cell state and a more differentiated sympathetic “adrenergic” cell state. These cell states are governed by autoregulatory transcriptional loops called core regulatory circuitries (CRCs), which drive the early development of sympathetic neuronal progenitors from migratory neural crest cells during embryogenesis. The adrenergic cell identity of neuroblastoma requires LMO1 as a transcriptional co-factor. Both LMO1 expression levels and the risk of developing neuroblastoma in children are associated with a single nucleotide polymorphism G/T that affects a <u>G</u>ATA motif in the first intron of LMO1. Here we show that wild-type zebrafish with the <u>G</u>ATA genotype develop adrenergic neuroblastoma, while knock-in of the protective <u>T</u>ATA allele at this locus reduces the penetrance of MYCN-driven tumors, which are restricted to the mesenchymal cell state. Whole genome sequencing of childhood neuroblastomas demonstrates that <u>T</u>ATA/<u>T</u>ATA tumors also exhibit a mesenchymal cell state and are low risk at diagnosis. Thus, conversion of the regulatory <u>G</u>ATA to a <u>T</u>ATA allele in the first intron of *LMO1* reduces the neuroblastoma initiation rate by preventing formation of the adrenergic cell state, a mechanism that is conserved over 400 million years of evolution separating zebrafish and humans.

## Introduction

Across all cell lineages and tissue types, groups of transcription factors act cooperatively in autoregulatory loops to regulate their own gene expression forming core regulatory circuitries (CRC), and gene expression across each circuit’s extended network, thereby governing cell identity and lineage specification (1). Arendt et al (2) have proposed that conservation of cell type identity across species reflects evolutionarily determined changes in the composition of the transcription factors that define key CRCs, leading orderly progression of conserved lineage-specific gene expression. Across vertebrates, a set of neural crest specific transcription factors facilitate the development of the neural crest, which gives rise to a wide spectrum of diverse cell lineages including neurons of the peripheral nervous system, glia, melanocytes, and facial bones and cartilage (3). In neuroblastoma, a pediatric malignancy originating from neural crest-derived progenitor cells, neuroblasts fail to differentiate and transform into a malignant cell state, often driven by aberrant high level expression of MYCN or MYC (4, 5). There are at least two primary CRC transcriptional networks in neuroblastoma that drive the growth and survival of neuroblastoma tumors. The most prevalent is the adrenergic CRC (including HAND2, GATA3, ISL1, PHOX2B, TBX2, and ASCL1), which is associated with committed progenitors of the sympathoadrenal cell lineage, while the mesenchymal CRC (including NOTCH2, BACH1, ID1, EGR3, FLI1, CBFB, and STAT3) represents a less differentiated mesenchymal or neural crest-cell-like transcriptional cell state (6–9). Interestingly, neuroblastoma tumors are often composed of both adrenergic and mesenchymal tumor cell populations that may spontaneously interconvert between networks, at least in part due to activation of the NOTCH pathway, which drives the mesenchymal cell state (10, 11). Therapeutic strategies are being designed to target the adrenergic and mesenchymal super-enhancer-associated transcriptional networks, with some success in model systems of neuroblastoma (7,10,12).

LIM-domain-only 1 (LMO1) acts an essential transcriptional co-regulator for the formation of the adrenergic neuroblastoma CRC (13), but is not part of the CRC itself because it is not a transcription factor and lacks a DNA binding domain. LMO1 facilitates transcription as a “bridge protein” by forming protein-protein interactions through its two zinc-finger LIM domains (14, 15). We have demonstrated that LMO1 overexpression synergizes with MYCN to accelerate tumor onset, penetrance and metastasis in a *dβh:MYCN*-driven zebrafish model (16).

Our genome-wide association study (GWAS) identified multiple single nucleotide polymorphisms (SNPs) in noncoding sequences across the genome associated with neuroblastoma susceptibility (17–20). In particular, one of the most significant neuroblastoma predisposition loci is the rs2168101 G>T transversion that resides within the first intron of the *LMO1* gene (20). In human neuroblastoma cells with the evolutionarily conserved G allele at this locus, the G creates the first position of a GATA-binding motif in the first intron of LMO1, which permits GATA3 to bind and leads to the formation of a large, tissue-specific super-enhancer that drives high levels of *LMO1* expression (20). By contrast, the T allele at this position forms the sequence TATA and prevents both GATA3 binding and formation of the intronic *LMO1* super-enhancer, leading to much lower levels of *LMO1* expression in neuroblasts and a lower risk of developing childhood neuroblastoma (20).

In our current study, we used genetic editing in the zebrafish to define the mechanism *in vivo* underlying the striking association between the germline rs2168101 G>T noncoding polymorphism in the first intron of *lmo1* and the risk of developing neuroblastoma. We found that the G allele is essential for the formation of the adrenergic CRC in human and zebrafish, so that individuals and fish with the T allele at this position only develop tumors relying on the mesenchymal CRC, which is much less efficient in initiating neuroblastoma.

## Results

### The rs2168101 locus is highly conserved throughout evolution

Previously, we showed that the human G>T polymorphism at the rs2168101 locus within the first intron of the *LMO1* gene is comprised of either a G, which represents the permissive allele associated with increased risk of developing neuroblastoma, or a T, which is protective of developing neuroblastoma (20). The evolutionary history of the G>T polymorphism is shown in Figure 1 A, which illustrates the remarkable finding that the G allele is exclusively found at this position throughout evolution except for human populations, which are the first to include individuals with the T allele. Interestingly, the T allele does not appear even in highly related non-human primates such as gorilla and orangutan (Supplementary Fig. S1). The reference G allele, by contrast, can be tracked back at least 400 million years to osteichthyes, the common ancestor of zebrafish and humans (Fig. 1 A).

**Figure 1:**
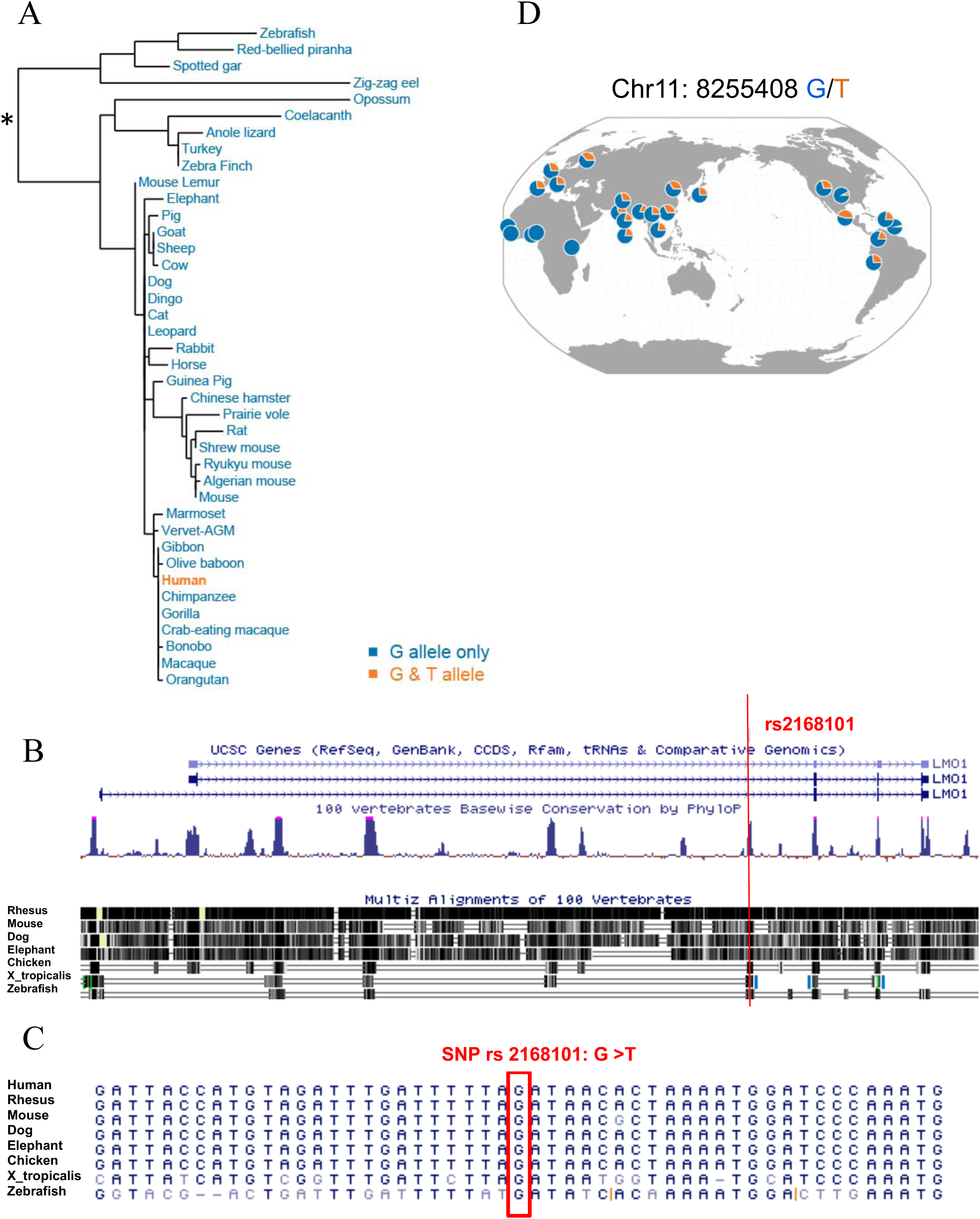
Evolutionary history of the G/T polymorphism at rs2168101. (A). Phylogenetic tree representing the evolutionary relationship between the *LMO1* genes from the indicated species over the last 400 million years. Blue font denotes species that exclusively harbor a G at rs2168101. Orange font denotes that only humans demonstrate a G/T polymorphism at the rs2168101 locus. (B). Distribution of the G and T alleles of rs2168101 across Human Genome Diversity Project (HGDP) populations, as illustrated by their genome browser (https://popgen.uchicago.edu). Circles create a pie chart in which blue represents the proportion of human populations from different parts of the world with a G at rs2168101 (human chromosome 11, position 8255408), and orange represents the proportion with a T. (C). Shown is a modified UCSC Genome Browser window of the human *LMO1* locus indicating the two alternative transcription start sites and the position of rs2168101 in the first intron (top), a vertebrate conservation track graphing PhyloP conservation scores (middle), and Multiz alignments of multiple vertebrate species (bottom), illustrating a high level of conservation in the non-coding region surrounding rs2168101. (D). The immediate sequence neighborhood surrounding rs2168101 in the first intron of *LMO1* from multiple species is shown. The G at rs2168101 in the human consensus sequence is marked with a red box.

Most noncoding intronic sequences within the *lmo1* gene are not well conserved between zebrafish and humans. The few exceptions are regions that consist of a few hundred base pairs of highly conserved sequence, including the region containing the rs2168101 locus, which likely contain important regulatory motifs (Fig. 1 B). We found that the noncoding region surrounding the conserved G allele at rs2168101 is highly conserved (Fig. 1 B and C), and 73% of the flanking 20 bp on each site of the G allele are identical between human and zebrafish (30 bp out of 41 bp) (Fig. 1 D). The *lmo1* coding sequence is also highly conserved among vertebrates (Fig. 1 C), with 84.5% (398/471) nucleotide identity between human and zebrafish and 98% identity at the amino acid level, including two perfectly identical LIM domains (Supplementary Fig. S4 D).

Interestingly, the T allele at rs2168101 in humans is well represented at comparable frequencies in the European, Asian, and American populations whereas it is nearly absent in Africans (Fig. 1 D, Supplementary Fig. S2), suggesting that the T allele arose in human populations as a single mutational event around the time of human migration out of Africa. These data in combination with our previous studies are consistent with a role for the G allele as part of a highly conserved regulatory element controlling *LMO1* expression in the developing PSNS(20).

### Substitution of a T allele for the G allele at rs2168101 impairs the initiation of neuroblastoma in a MYCN-driven zebrafish model

To dissect the mechanism through which the T allele at rs2168101 protects children from developing neuroblastoma, we used transcription activator-like effector nuclease (TALEN)-mediated gene editing to generate the rs2168101 T allele in zebrafish (Fig. 2 A and Supplementary Fig. S3; see Methods for details). Because the G at rs2168101 comprises the first nucleotide of a highly conserved *GATA* site, we designated this heterozygous mutant line “*lmo1 GATA/TATA”* (also referred to herein as the “*GATA/TATA”* line). We found that the heterozygous *GATA/TATA* and homozygous *TATA/TATA* fish were viable, developmentally normal, and fertile. We crossed the *GATA/TATA* line to our previously described *Tg(dβh:MYCN;dβh:EGFP)* transgenic zebrafish line in which *MYCN* and *EGFP* expression are driven in the PSNS by the dopamine beta-hydroxylase promoter and *MYCN* overexpression gives rise to neuroblastoma in the inter-renal gland (IRG) (Fig. 2 B and 2 C(21). The IRG is the zebrafish equivalent of the adrenal medulla, which is the most frequent site of human neuroblastoma (4, 21). We have shown that the tumors that arise in the IRG in this transgenic fish model are small, round blue cell tumors that express tyrosine hydroxylase and synaptophysin, which are markers of human neuroblastoma (21).

**Figure 2:**
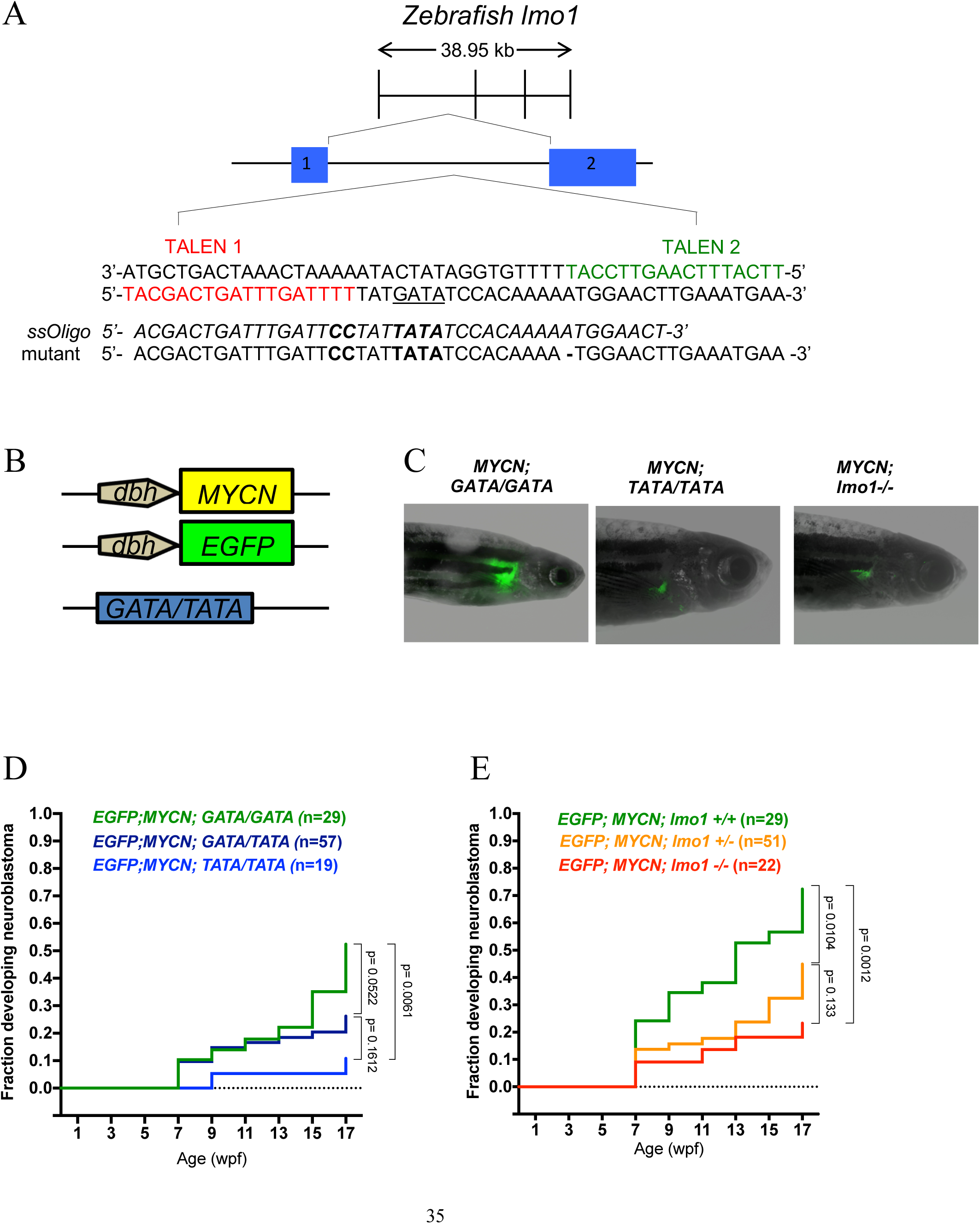
The development of *MYCN*-driven neuroblastomas is impaired in the *TATA/TATA* and *lmo1*-null backgrounds. (A). Diagram illustrating the construction of the *lmo1 GATA/TATA* (*GATA/TATA*) zebrafish line in which TALEN-mediated gene editing was used to replace the G at rs2168101 with a T. The rs2168101 G resides within the first intron of the zebrafish *lmo1* gene (exons 1 and 2 are denoted by blue boxes) and creates the first nucleotide of a *GATA* DNA-binding sequence (in bold). To facilitate the precise genome editing and knock-in of the T allele at this locus, we used TALEN gene-editing technology targeting the sequences flanking the G at rs2168101 (as indicated in red and green) together with a single-stranded DNA oligonucleotide containing a T instead of the G with short flanking homology arms of 20 nucleotides (*TATA-ssOligo*). To prevent TALEN binding to the 5′ arm and activity after successful knock-in of the *TATA-ssOligo* and to aid in the identification of embryos containing the modified sequence, the *TATA-ssOligo* was designed with two additional nucleotide changes (CC to replace TT in the 5’ homology arm, marked in bold) to create a new restriction site for TfiI (see also Fig. S3). (B). To analyze the effect of the rs2168101 G>T substitution on *MYCN*-induced neuroblastoma, compound transgenic zebrafish lines were created by crossing the transgenic lines *Tg(dβh:MYCN) and Tg(dβh:EGFP)* with the *GATA/TATA* knock-in line, as illustrated. The *dβh:EGFP* and *dβh:MYCN* lines, in which the zebrafish *dβh* promoter was used to facilitate tissue-specific expression of *EGFP and MYCN,* were established previously(21). (C). Representative fluorescent images of adult zebrafish showing EGFP-expressing tumors arising in the indicated transgenic lines. (D-E). Starting at 5 wpf, zebrafish with the indicated genotypes were monitored biweekly for the presence of tumors by EGFP fluorescence microscopy. The graph shows a Kaplan-Meier analysis of the cumulative frequency of neuroblastomas in the transgenic lines. Statistical analysis was performed using the logrank test.

Three different genetically modified zebrafish lines were generated to determine the influence of the *TATA/GATA* regulatory site on the rate of initiation and penetrance of neuroblastoma: i) *Tg(dβh:MYCN; dβh:EGFP; GATA/GATA*), ii) *Tg(dβh:MYCN; dβh:EGFP; GATA/TATA*), and iii) *Tg(dβh:MYCN; dβh:EGFP; TATA/TATA*). We monitored offspring for the onset of EGFP+ tumor masses in the anterior region of the abdomen where the IRG is located (Fig. 2 C). Although tumors arose more frequently in the GATA/GATA than in the GATA/TATA genotype (Fig. 2 D), the overall tumor onset curves for *GATA/GATA* fish and *GATA/TATA* fish were not statistically significantly different, suggesting that in this model one intact GATA site was sufficient to promote neuroblastoma. By contrast, only 10% of the *TATA/TATA* fish developed neuroblastoma over the first 17 weeks of life, which represented a significantly lower penetrance than the tumor onset for the *GATA/GATA* fish (*p*<0.01). This finding is consistent with the significant overrepresentation of homozygous *GATA/GATA* genotypes in neuroblastoma revealed by GWAS in children, in that 57.9% of children with neuroblastoma had GATA/GATA, compared with 35.6% for *GATA/TATA*, and only 6.4% for the *TATA/TATA* genotype (20). Thus, our studies in the zebrafish model are very consistent with GWAS findings in children demonstrating that the G allele at rs2168101 increases the risk of developing neuroblastoma while the T allele at this position is protective.

### Knockout of lmo1 in zebrafish reduces the penetrance of MYCN-induced neuroblastoma

To independently test whether complete loss of *lmo1* expression confers protection against the development of neuroblastoma, we used CRISPR-Cas9-mediated gene editing to generate a *lmo1* knockout allele containing a 32-basepair deletion of coding sequences in the second exon that led to premature termination of translation (Supplementary Fig. S4). As previously reported in an *Lmo1* mouse knockout model (22), we found that zebrafish with homozygous knockout of the *lmo1* gene were viable, developmentally normal and fertile. The *lmo1+/-* line was bred to the *Tg(dβh:MYCN; dβh:EGFP)* line to generate zebrafish with the following genotypes for analysis: *Tg(dβh:MYCN; dβh:EGFP; lmo1+/+*), *Tg(dβh:MYCN; dβh:EGFP; lmo1+/-*), and *Tg(dβh:MYCN; dβh:EGFP; lmo1-/-*). Analysis of these three lines (Fig. 2 E) showed a significantly reduced penetrance of neuroblastoma at 17 weeks of life in *lmo1*-/- compared to *lmo1+/+* zebrafish (22% versus 70% respectively; *p*<0.01). The heterozygous *lmo1+/-* line showed an intermediate tumor penetrance with 41% of fish developing tumors at 17 weeks. Thus, the protective effect of the G>T substitution at rs2168101 (Fig. 2 D) was similar to the effect of complete loss of expression of a functional *lmo1* gene (Fig. 2 E). This indicates that elevated *lmo1* expression in neuronal progenitor cells accounts for the increase in neuroblastoma penetrance associated with the G allele at rs2168101.

### Other Lim-only protein family members generally don’t compensate for diminished lmo1 expression in zebrafish MYCN-driven neuroblastomas

*LMO1, LMO2* and *LMO3* are functionally redundant T-ALL oncogenes (23–25), so we reasoned that loss of *lmo1* expression in zebrafish might stimulate the upregulation of other *lmo* family members during neuroblastoma pathogenesis. We therefore analyzed gene expression in *MYCN*-induced neuroblastomas in *GATA/GATA*, *TATA/TATA* and *lmo1*-/- genetic backgrounds by RNA-Sequencing (RNA-Seq). Consistent with human neuroblastomas (20), *lmo1* mRNA expression levels in neuroblastomas from *GATA/GATA* fish were significantly higher than in tumors from either *TATA/TATA* or *lmo1-/-* fish. The *lmo1-/-* fish were included in this study as a control group of fish that lack intact lmo1 protein expression, and the lower lmo1 mRNA levels in these fish are likely due to nonsense-mediated degradation of the mutant lmo1 mRNA containing a premature stop codon in exon 2 (Fig. 3 A and Supplementary Fig. S4). Importantly, expression of the other six highly related zebrafish *lmo* family members (*lmo2, lmo3, lmo4a, lmo4b, lmo5, lmo6, lmo7a* and *lmo7b*) was much lower than *lmo1* in the *GATA/GATA* fish and showed no appreciable differences between the three tumor genotypes (Fig. 3 A). Thus, there is no obvious compensatory overexpression of other *lmo* family members due to the low levels of *lmo1* mRNA expression in zebrafish *TATA/TATA* tumors.

**Figure 3:**
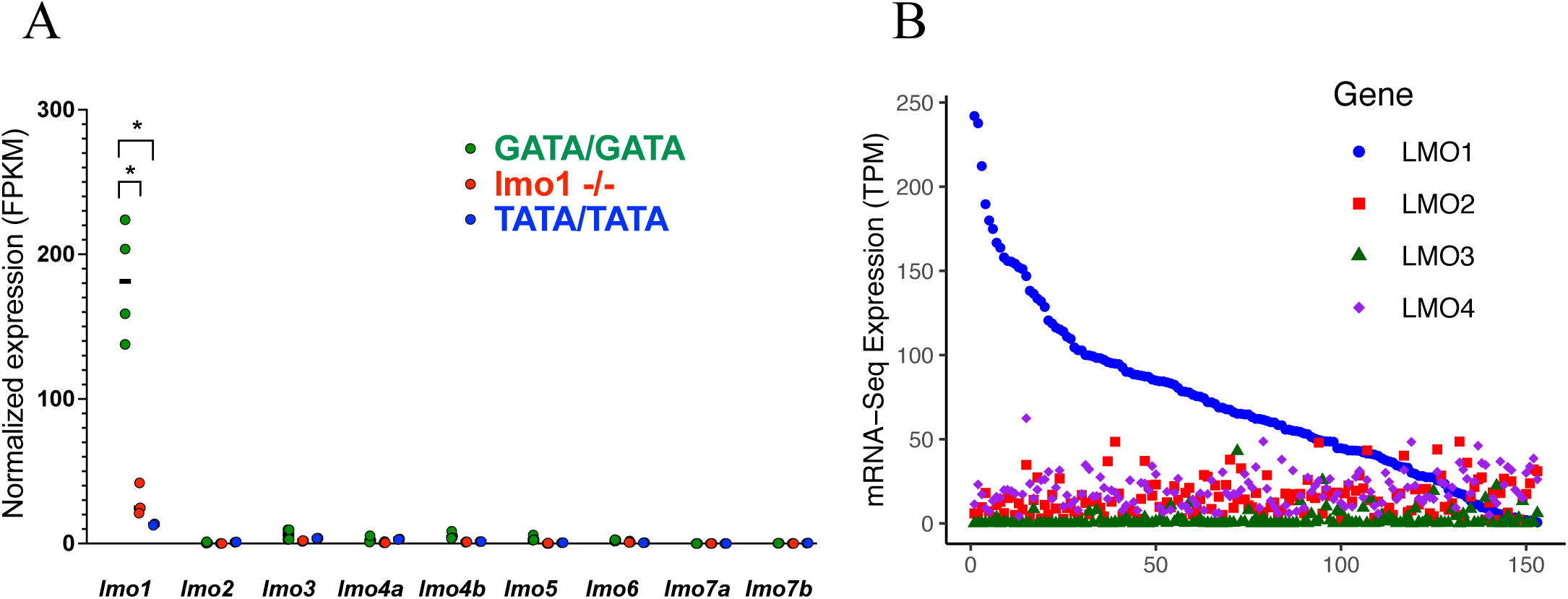
Expression of the *lmo* family genes in zebrafish and human neuroblastoma. a. RNA-Seq analysis was performed to measure the relative mRNA expression of *lmo* family genes in neuroblastomas arising in zebrafish with the indicated genotypes (*GATA/GATA*, n = 4; *lmo1-/-*, n = 3; and *TATA/TATA*, n = 2). mRNA expression levels for the indicated *lmo* family genes are represented by FPKM log-scale values. Expression < 1 FPKM is considered as non-expressed. Statistical analysis was performed using the two-tailed, unpaired *t* test. \**p* < 0.005. b. Relative *LMO1-4* mRNA expression levels measured by RNA-Seq in 153 primary human neuroblastoma samples (from TARGET), ranked from highest (left) to lowest levels of *LMO1* expression (in FPKM). Expression correlation analysis demonstrated weak inverse correlation between LMO1 and the other three LMO Family member. (R>= -0.3, p<0.05)

Similarly, analysis of 153 primary human neuroblastoma samples by RNA-Seq showed a weak inverse correlation between *LMO1* expression and the expression levels of the other *LMO* family members (R>= -0.3, p<0.05) (Fig. 3 B). The weakness of this correlation is illustrated by the fact that the strongest inverse correlation was observed between *LMO1* and *LMO3* expression levels (R= -0.3), meaning that only about 10% (R^2^ ∼ 0.1) of the *LMO3* expression level can be explained by low *LMO1* expression in human neuroblastoma samples. Since LMO1 has a central role as a transcriptional cofactor in the adrenergic neuroblastoma core regulatory circuitry (13), the lack of compensation by other *lmo* family members in zebrafish tumors with low *lmo1* expression in *MYCN*-driven neuroblastomas suggests that neuroblastomas arising in fish with the *TATA/TATA* and *lmo1-/-* genotypes employ mechanisms of transformation independent of lmo-family-proteins.

### Lmo1 co-regulates the adrenergic neuroblastoma CRC

LIM-only proteins are well known to function as “linker” proteins that facilitate the assembly of transcription factor complexes involved in tissue-specific gene regulation (26–29). We have shown that LMO1 functions in this capacity as an essential transcriptional cofactor for the human adrenergic neuroblastoma CRC (13). To analyze the neuroblastoma CRC in *MYCN*-driven tumors with disrupted *lmo1* expression, we used RNA-Seq to compare the transcriptomes of zebrafish *MYCN*-driven neuroblastomas from the *GATA/GATA*, *TATA/TATA* and *lmo1*-/- lines. To study the gene expression of EGFP-expressing neuroblastoma cells by RNA-Seq analysis, we dissected the EGFP-fluorescent tumors from juvenile zebrafish, mechanically prepared a single cell suspension and then sorted the EGFP+ cells prior to RNA extraction. Thus, our total RNA-seq analysis focuses on MYCN- and EGFP-overexpressing neuroblastoma cells and does not address changes in gene expression of the surrounding stroma cells.

Figure 4 A provides an overview of significant differences in gene expression of zebrafish tumor cells in the *GATA/GATA*, *TATA/TATA* or the *lmo1*-/- genetic backgrounds (*p*<0.05 based on an absolute log2 fold change compared to GATA/GATA of >= 0.45). Ninety transcription factor genes were significantly lower expressed in neuroblastomas from both *TATA/TATA* and *lmo1*-/- zebrafish compared to tumors in the GATA/GATA background (Fig. 4 B). These differentially expressed transcription factors include *lmo1* as well as *gata3*, *hand2*, *phox2b*, *isl1*, and *ascl1*, which form the adrenergic CRC of neuroblastoma (Fig. 4 C) (7, 13). In addition, *LMO1* expression levels are also higher in human neuroblastoma cell lines of the adrenergic subtype and lower in the mesenchymal type (Fig. 4 D), consistent with our recent study showing that LMO1 is an essential transcriptional cofactor of the adrenergic CRC (13). These results indicate that high levels of *lmo1* expression by the neuroblastoma cells are required to establish the adrenergic cell state.

**Figure 4:**
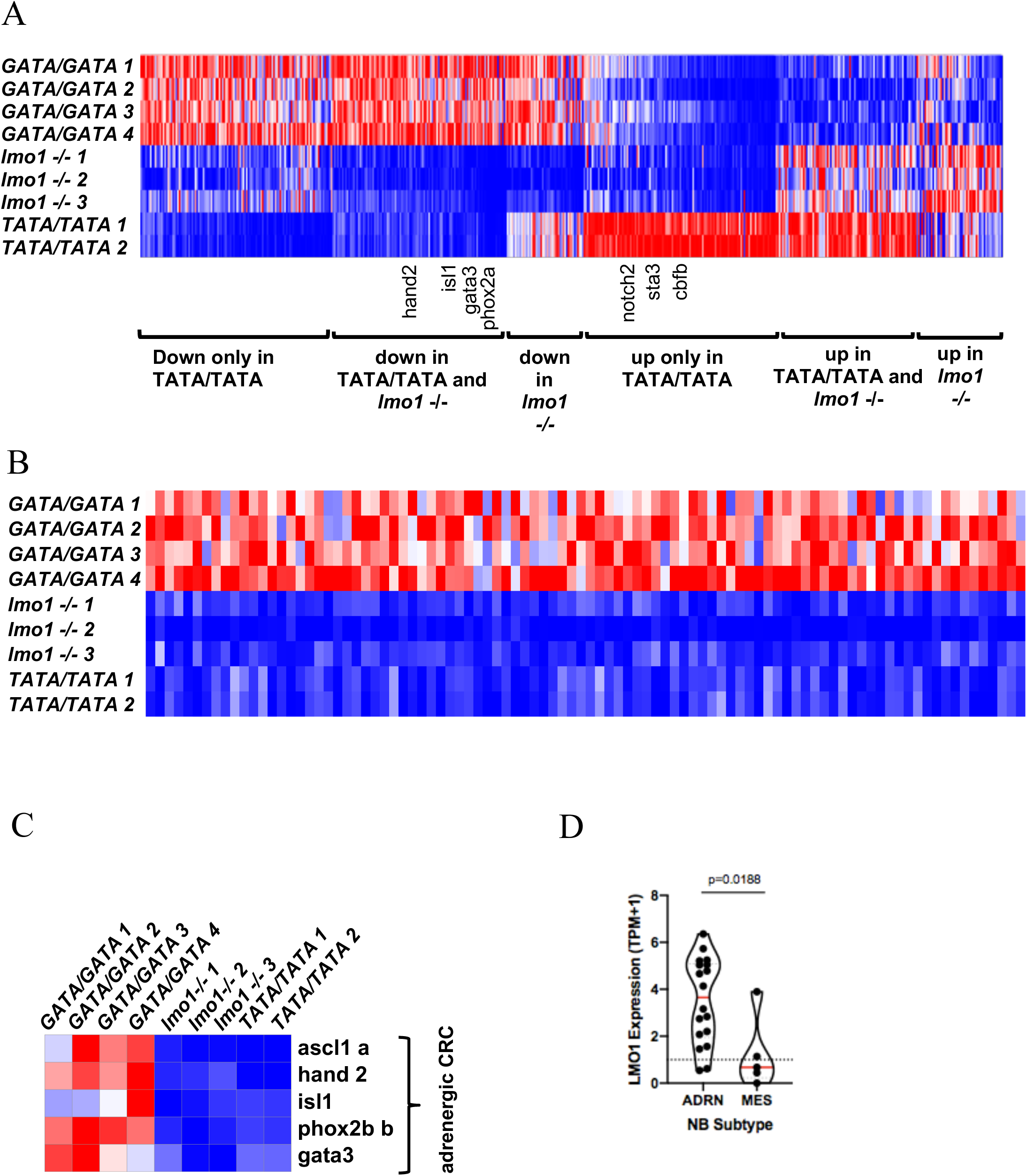
*Lmo1* co-regulates transcription factors that comprise the adrenergic neuroblastoma CRC. (A). Heatmap image based on RNA-Seq data analysis showing differentially expressed genes in *MYCN*-induced neuroblastoma tumors arising in *lmo1 GATA/GATA (*wild-type), *lmo1*-/- and *lmo1 TATA/TATA* backgrounds categorized into 6 groups as indicated. Each row corresponds to a gene, and signal intensity is normalized across the row. Genes were rank ordered from highest (right side of the map) to lowest (left side of the map) based on fold change of gene expression in *TATA/TATA* or *lmo1-/-* compared to *GATA/GATA*. (B). Heatmap of transcription factors that are downregulated in *MYCN*-induced neuroblastomas arising in both *lmo1*-/- and *TATA/TATA* backgrounds compared to *GATA/GATA.* Each row corresponds to a transcription factor gene, and signal intensity is normalized across the row. Genes were rank-ordered based on fold-change values between *GATA/GATA* and *TATA/TATA* tumors from highest (right side of the map) to lowest (left side of the map). (C). Heatmap representing gene expression changes of the known adrenergic neuroblastoma CRC transcription factors *isl1, gata3, ascl1, phox2b,* and *hand2* in *MYCN*-induced neuroblastoma tumors arising in the *GATA/GATA, lmo1-/-* and *TATA/TATA* backgrounds. (D). *LMO1* mRNA expression (TPM+1) violin plots retrieved from the 21Q1 release of Depmap (^18^), from 19 neuroblastoma cell lines. Cell lines were defined as adrenergic (ADRN) (n=14) or mesenchymal (MES) (n=5) subtypes based on general gene expression profiles. Red bars indicate mean, dotted line indicates a TPM+1 of 1. p=0.0188 by student’s t-test.

### Zebrafish TATA/TATA neuroblastomas adopt the mesenchymal CRC

Because neither *lmo1-/-* nor *TATA/TATA MYCN*-expressing tumors appeared to be driven by the adrenergic CRC, which is preferred by neuroblastomas arising in the GATA/GATA background, we questioned whether either tumor type was instead driven by the mesenchymal CRC (6,7,10). Further analysis of the RNA-Seq data revealed that mesenchymal neuroblastoma CRC transcription factors (including *notch2*, *id1*, *egr3*, *irf3*, *cbfb, bach1a, bach1b, tcf7l2,* and *mef2b)* (6) were upregulated in the *TATA/TATA* tumors, compared to *GATA/GATA*, but not in the *lmo1-/-* tumors (Fig. 5 A). Consistent with these findings, *TATA/TATA* tumors also showed upregulation of the mesenchymal marker fibronectin 1a (*fn1a;* Fig. 5 B) (6).

**Figure 5:**
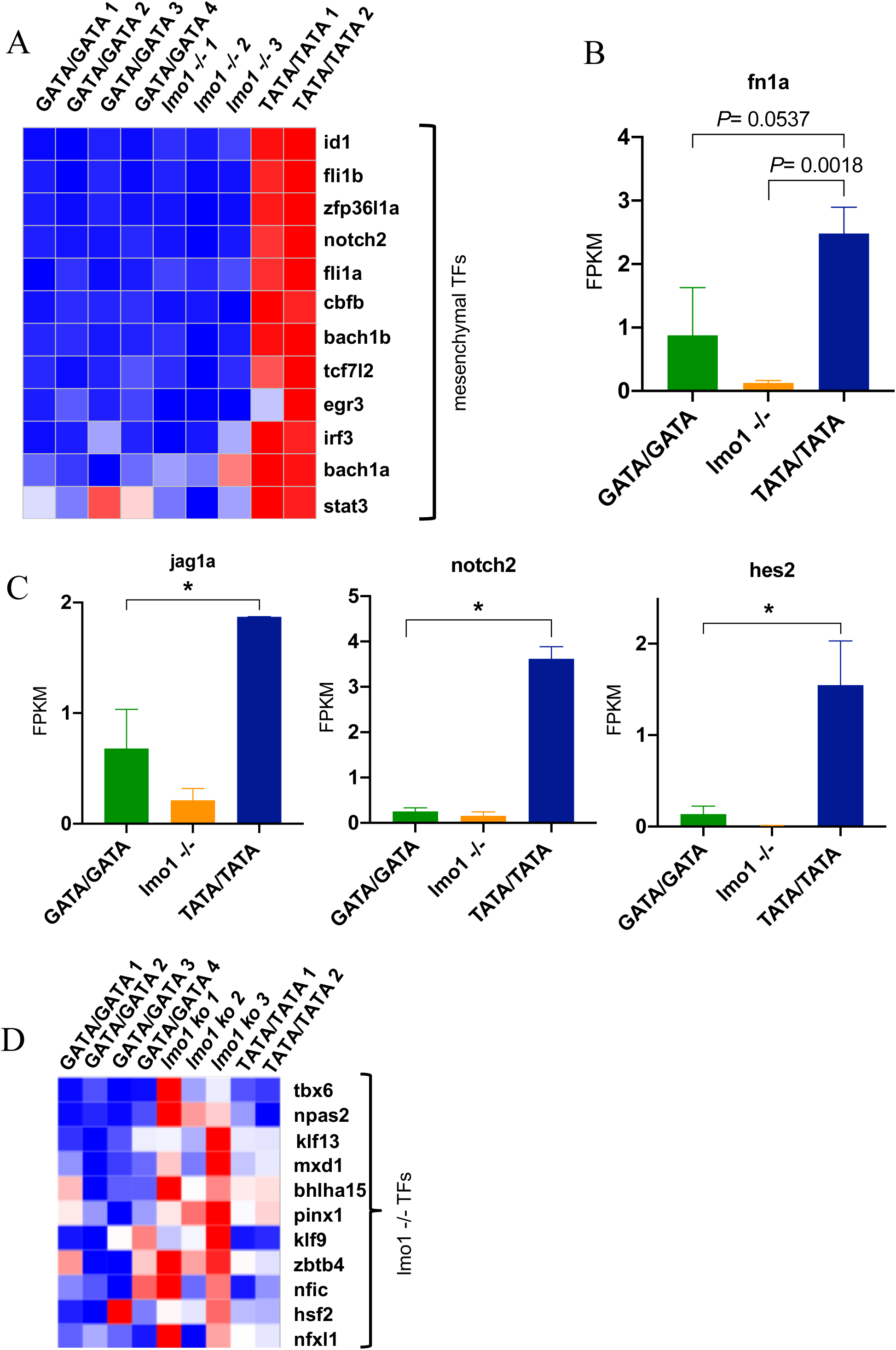
*MYCN*-driven neuroblastomas from the *TATA/TATA* line, but not the *lmo1-/-* line, express the mesenchymal CRC. (A). Heatmap image based on RNA-Seq data analysis showing the indicated genes of the mesenchymal CRC, which are upregulated in the *MYCN*-induced neuroblastomas arising in the *TATA/TATA* background, but not in the *lmo1*-/- background, when compared to the *GATA/GATA* zebrafish. (B). Relative mRNA expression levels of the mesenchymal neuroblastoma phenotype marker *fn1a* in neuroblastoma arising in zebrafish with the indicated genotypes. (C). Relative mRNA expression levels of NOTCH pathway genes, including *jag1a*, *notch2*, and *hes2*, in neuroblastomas arising in zebrafish from the indicated genotypes. Statistical analysis was performed using the two-tailed, unpaired *t*-test. * *p* <= 0.003. (D). Heatmap of transcription factors exclusively upregulated in *MYCN*-driven neuroblastomas from the *lmo1*-/- line. TFs; transcription factors.

Because the NOTCH pathway was shown to reprogram adrenergic neuroblastoma cells to adopt a mesenchymal cell state (10), we examined the RNA-Seq data for genes associated NOTCH pathway upregulation. Indeed, *TATA/TATA* tumors, but not *lmo1-*/- tumors, showed upregulation of notch receptor (*notch2b*) and ligand (*jagged1*) genes, as well as the NOTCH target gene *hes2* (Fig. 5 C). Interestingly, the *lmo1-/-* tumors did not highly express the signature genes from either the adrenergic or mesenchymal cell states (Fig. 4 D and 5 A), but instead expressed an alternative set of transcription factors with no previously defined role in either of these two neuroblastoma CRCs (Fig. 5 D).

### Human TATA/TATA neuroblastomas resemble zebrafish TATA/TATA tumors with low LMO1 expression and a mesenchymal expression profile

To test the hypothesis that human neuroblastomas arising in the *TATA/TATA* background have low *LMO1* expression and, as in the zebrafish model, are enriched for neuroblastomas relying on the mesenchymal rather than the adrenergic CRC, we used a new publicly available data set from the Gabriella Miller Kids First Data Research Program. This data set includes 124 tumors from low-, intermediate- and high-risk neuroblastoma cases (30), for which both rs2168101 genotypes and tumor cell RNA-Seq results are available. There were seven neuroblastomas with *TATA/TATA* genotypes, which showed significantly lower *LMO1* expression levels than *GATA/TATA* or *GATA/GATA* tumors (p<0.005; Fig. 6 A). It is notable that six of the seven TATA/TATA tumors were classified as low risk (Fig. 6 A). Previously published results were based on the TARGET neuroblastoma dataset, which only included high-risk neuroblastoma patients, and none of the neuroblastomas in the TARGET dataset had the TATA/TATA genotype (20).

**Figure 6:**
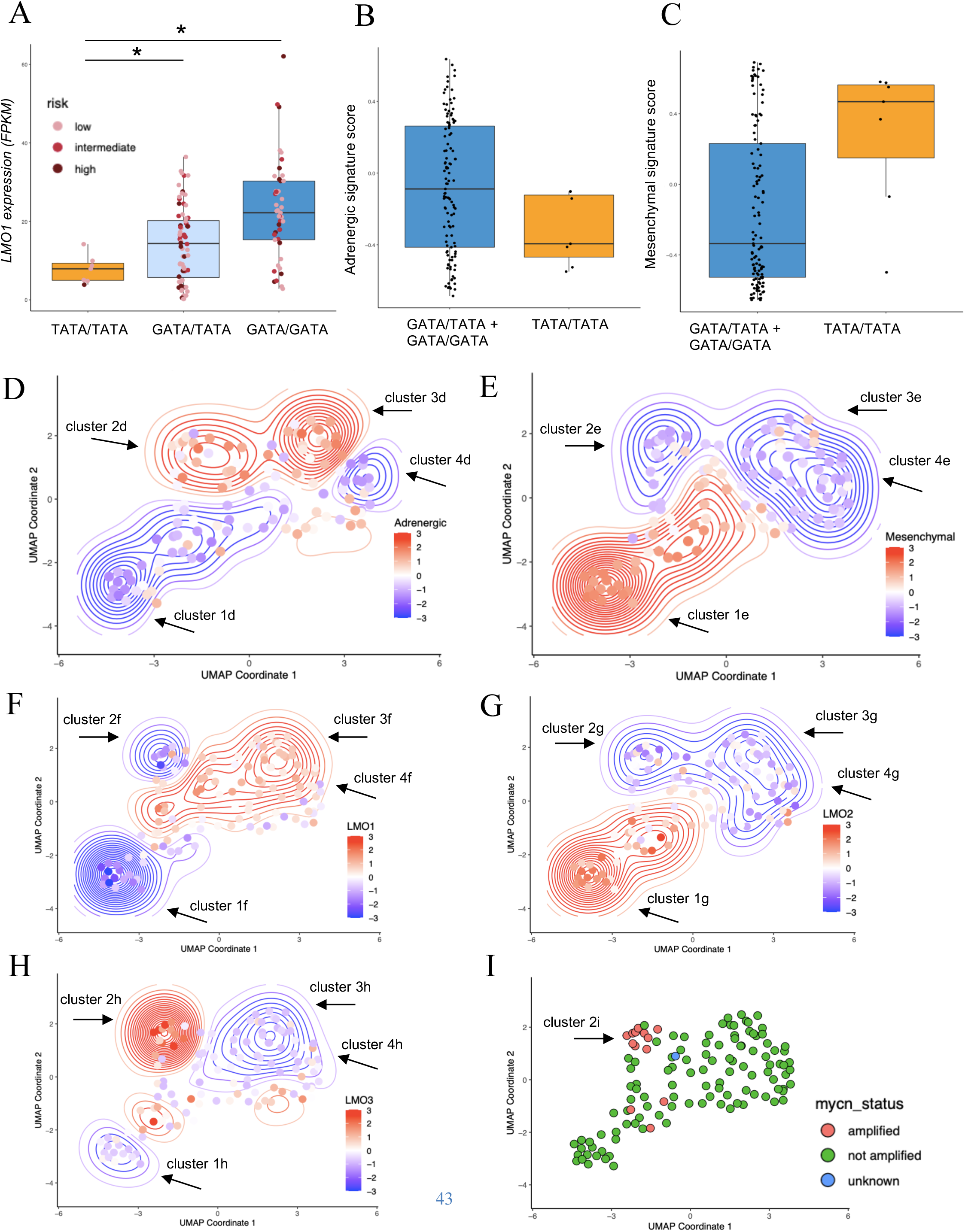
Human *TATA/TATA* neuroblastomas resemble zebrafish *TATA/TATA* tumors with low *LMO1* expression and a mesenchymal phenotype. (A). Relative LMO1 mRNA expression levels (in FPKM) in 124 human neuroblastoma samples with the indicated genotypes at rs2168101: TATA/TATA, GATA/TATA or GATA/GATA. Each neuroblastoma sample is assigned to either low (pink), intermediate (light red) or high risk (purple). Statistical analysis was performed using the two-tailed, Welch’s *t* test. \**p* < 0.005. (B-C). Adrenergic and mesenchymal gene set signature scores were generated using Gene Set Variation Analysis (GSVA) based on the previously published expression profiles by Von Groningen et al. (*8*). Higher positive scores indicate upregulation of the corresponding signature whereas lower negative scores indicate down regulation. GATA/GATA and GATA/TATA tumors were combined into one group (blue) and compared to the TATA/TATA tumors (tan). (D-E). Uniform manifold approximation and projection (UMAP) representing the whole transcriptome landscape of 124 human neuroblastoma samples (dots) in a two-dimensional space combined with the adrenergic (D) and mesenchymal (E) gene signatures. Relative Z-score-transformed expression for each signature (according to the heat scale) is shown for each tumor (points) and overall density (contours). Clusters 1-4 represent tumors falling into similar density contours based on the adrenergic signature (D) or mesenchymal signature (E). (F-H). Combination of UMAP dimensionality reduction analysis and expression levels of *LMO1* (F), *LMO2* (G) and *LMO3* (H). Relative Z-score of log2-transformed expression for *LMO1* (F), *LMO2* (G) or *LMO3* (H) (according to the heat scale) is shown for each tumor (points) and overall density (contours). (I). MYCN status for all 124 human neuroblastoma samples.

We next asked whether the TATA/TATA genotype was significantly associated with usage of the mesenchymal CRC in human neuroblastoma, as we observed in zebrafish neuroblastoma (Fig. 5). For this analysis we applied a gene set signature score for every patient using adrenergic and mesenchymal signatures (6) and performed Gene Set Variation Analysis (GSVA). As in the zebrafish, we observed that none of the seven human neuroblastomas with the *TATA/TATA* genotype had a positive adrenergic CRC signature score (Fig. 6 B). By contrast, neuroblastomas with *GATA/GATA* and *GATA/TATA* genotypes were approximately equally divided between those with predominately adrenergic and predominantly mesenchymal CRC signature scores (Fig. 6 B and C). This is consistent with known plasticity within human neuroblastomas, such that tumor cells can shuttle back and forth between these signatures based, at least in part, on the levels of NOTCH signaling (10). Within the *TATA/TATA* human tumors the majority exhibited a positive score for the mesenchymal signature, which is consistent with our results in the zebrafish model (Fig. 5 A).

### Visualization of patterns of gene expression in primary human neuroblastomas

To help visualize different subsets of neuroblastomas, we employed Uniform Manifold Approximation and Projection (UMAP) dimensionality reduction analysis based on the gene expression values for each tumor, and then investigated how other features varied across the two-dimensional UMAP embedding. We first analyzed the adrenergic signature score in the context of the UMAP values, which revealed four clusters of neuroblastomas (Fig. 6 D). Clusters 1d and 4d represent tumors with largely downregulated adrenergic signature scores (blue colored density contours), and clusters 2d and 3d with largely upregulated adrenergic signature scores (red colored density contours) (see Fig. 6 D). By contrast, the mesenchymal signature in Figure 6 E is represented largely by a reciprocal pattern, in that it is upregulated within tumors mapped by contours corresponding to cluster 1e in Figure 6 E, while it is downregulated in tumors falling within the contours of clusters 2e to 4e. Interestingly, cohort 4d tumors not only have low adrenergic signature scores, but also have low mesenchymal signature scores, possibly indicating that they are driven by an alternative CRC with a different signature (31).

In Figure 6 F, showing the UMAP coordinates grouped by *LMO1* expression levels, the tumors in cluster 1f with low *LMO1* expression levels correspond to a subset of the tumors with low adrenergic scores (cluster 1d; Fig. 6 D) and high mesenchymal scores (cluster 1e; Fig. 6 E). This cluster contains six of the seven TATA/TATA tumors, which is consistent with their low *LMO1* expression levels. Interestingly, tumors within cluster 2f in Figure 6 F also have low *LMO1* expression levels but can adopt the adrenergic signature (cluster 1d; Fig. 6 D). To determine whether this apparent discrepancy might be due to high levels of expression of *LMO* family members other than *LMO1*, we assessed *LMO2* and *LMO3* expression levels across the UMAP coordinates (Fig. 6 G and H). *LMO2* is very low overall as shown in Figure 3 B, and except for rare cases with moderate *LMO2* expression, the contours representing *LMO2* expression in Figure 6 G do not reach levels at which *LMO2* could substitute for *LMO1*, consistent with other studies in mice implicating *lmo1* and *lmo3* as the closely related LMO family members expressed normally by neuronal cells (22). Higher levels of *LMO3* expression occurred in some of the human neuroblastomas corresponding to cluster 2h based on the adrenergic score (Fig. 6 H), explaining how the adrenergic signature can form in these tumors despite relatively low levels of *LMO1* expression. Interestingly, the human neuroblastomas expressing high levels of *LMO3* in cluster 2h are enriched for neuroblastomas with *MYCN* gene amplification (Fig. 6 I, cluster 2i). Tumors mapped mainly to cluster 3h have high *LMO1* expression levels (cluster 3f; Fig. 6 F) and have high adrenergic signature scores and low mesenchymal signature scores (cluster 3d and 3e in Fig. 6 D and E).

Thus, detailed gene expression studies in both zebrafish and human neuroblastomas reveal conservation of tumor-cell-specific gene expression signatures driven by the adrenergic and mesenchymal CRCs. The role of LMO1 as an essential cofactor for the adrenergic subtype of neuroblastoma appears to have been highly conserved throughout the 400 million years of evolution that separate zebrafish and human populations.

## Discussion

Many SNPs tightly linked by GWAS to disease phenotypes are located within non-coding regions of the genome, implying that they affect regulatory sequences and alter the expression of genes rather than the structure of the encoded proteins (32–35). Functional *in vivo* analysis of SNPs implicated by GWAS so far has generally not included the introduction of the orthologous base change in the noncoding genome, but rather has mainly focused on *in vitro* reporter assays of gene expression, epigenetic analysis of the associated gene locus and *in vivo* knockout models of the implicated gene (36–38). However, studying the precise regulatory SNPs in animal models, as we have done here, is of critical importance to identify the causal mechanisms and context that regulate the expression levels of disease-associated genes. One example of the usefulness of *in vivo* models for study of SNPs implicated by GWAS is the work of Madelaine and coworkers (39). Their work shows that deletion of highly conserved non-coding DNA sequences, including a SNP linked to retinal vasculature defects, did not result in downregulation of MEF2C, the original candidate gene associated with this disease. Instead, the deletion of this region caused down-regulation of expression of the miRNA-9 gene, and programmed depletion of this microRNA reproduced the retinal vasculature phenotype (39). Thus, the use of genomic studies in the zebrafish animal model have helped to clarify mechanisms underlying genetic loci implicated by GWAS (40–42).

### The regulatory SNP rs2168101 predicts neuroblastoma risk in zebrafish as well as human populations

In our current study, we studied the impact and mechanism of rs216810, one of the most significantly associated SNPs with the risk of developing neuroblastoma in children. We first noted that wild-type zebrafish only have the G allele at the nucleotide corresponding to the G/T SNP at rs2168101 in human populations. This is expected because the G allele is conserved throughout vertebrate evolution, and the T allele polymorphism at this position arose first in human populations (Fig. 1). Hence, we used genome editing technology to introduce the T allele into the zebrafish germline and then bred this allele into our *MYCN* driven neuroblastoma model. We found significantly reduced penetrance of MYCN-driven neuroblastoma in transgenic zebrafish with the T/T allele at the rs2168101 locus (Fig. 2), concurrent with decreased *lmo1* expression by these tumors (Fig. 3). These results support our earlier work in human neuroblastoma cell lines establishing that the substitution of a T allele for the G allele disrupted a key GATA site and downregulated LMO1 expression by blocking the formation of a lineage-specific enhancer in the first intron of the *LMO1* gene (20). Our studies in the zebrafish model therefore support a causative role for the G allele at rs2168101 in upregulating *lmo1* expression levels in developing neuroblasts, which increases the rate of initiation of *MYCN*-driven neuroblastoma in this model (20).

### The CRC itself determines the frequency and penetrance of neuroblastoma

Studies by several laboratories have identified at least two predominate core regulatory circuits (CRCs) in childhood neuroblastoma, which are called the adrenergic and mesenchymal CRCs (6,7,10). The central role and conservation of tissue-specific CRCs in development and cancer is supported by our finding that neuroblastomas arising in wild-type GATA/GATA fish over-express genes typical of the adrenergic CRC (*hand2*, *gata3*, *isl1*, *ascl1* and *phox2a*; Fig 4D). In fish, as in human neuroblastomas, this permissive SNP drives high levels of expression LMO1, which is an essential transcriptional cofactor of the adrenergic CRC (13). By contrast, fish with the TATA/TATA allele specifically downregulate adrenergic CRC genes and instead exhibit high levels of expression of genes associated with the mesenchymal CRC, such as *notch2, id1, stat3, egr3, phox1, fli1,* and *mef2b (*Fig. 5a). Analogous to human neuroblastomas, the mesenchymal subtype of neuroblastoma in TATA/TATA zebrafish neuroblastomas is associated with activation of *hes2, jag1* and *notch2,* indicating activation of the NOTCH signaling pathway (Fig. 5c). This finding agrees with work of van Groningen and coworkers showing that activation of NOTCH signaling participates in the induction of the mesenchymal cell state in human neuroblastomas (10). In our study, we used cell sorting based on *dβh*-driven EGFP expression to direct our RNAseq studies to the dopamine B hydroxylase-expressing, EGFP+ neuroblastoma cells. This has the advantage that it focuses the analysis on the tumor cell population in these fish but has the disadvantage that it excludes stromal cells of the tumor niche and infiltrating blood cell populations, which may collaborate in unique ways in the tumor environments that develop in individuals with different germline genotypes. Future studies are warranted to capitalize on newer single cell RNAseq technologies of unsorted cells from the tumors in these fish, which will permit simultaneous analysis of the gene expression not only by EGFP-positive cells, but also will include gene expression by the many types of cells in the tumor niche that participate in the formation of MYCN-driven neuroblastoma.

### The TATA/TATA genotype at rs2168101 favors the onset of neuroblastomas with mesenchymal rather than adrenergic CRCs

Our current study also includes new information from a new genome DNA and RNA sequencing dataset by 124 primary childhood neuroblastomas, created with support of the Gabriella Miller Kid’s First Pediatric Research Program (Fig. 6). This new dataset is particularly important because in earlier studies we used the TARGET dataset, which was limited to tumors from children with high-risk neuroblastoma. Our new analysis shows neuroblastomas arising in the *TATA/TATA* background generally are more localized and lower risk, and that these tumors do not adopt the adrenergic gene expression program, but rather often use the mesenchymal CRC (Fig. 6b and c).

Our finding that the relatively infrequent tumors that arise in the TATA/TATA zebrafish exhibit the mesenchymal CRC signature is consistent with our previous studies showing that a high level of LMO1 expression is required for the adrenergic CRC to form in human neuroblastoma (13). In agreement with these findings in the TATA/TATA zebrafish, our new data using the Gabriella Miller human neuroblastoma dataset also shows that human tumors arising in children with the TATA/TATA genotype at rs2168101 exhibit the mesenchymal signature, and most of these tumors are assigned to children with low-risk, localized neuroblastomas that are often cured by surgery alone (Fig.6). An apparent paradox emerges, however, because others have shown that GATA/GATA or GATA/TATA neuroblastoma may shift from the adrenergic to the mesenchymal cell state under the selection pressure of chemotherapy, presumably because the mesenchymal cell state imparts drug resistance (6, 43). However, this represents an apparent paradox, because the mesenchymal cell state appears to be associated both with the initiation of localized tumors in young children and with rapidly growing and metastatic drug resistant neuroblastomas that are not responding to therapy. To explain this apparent discrepancy, we hypothesize that adrenergic neuroblastoma cells can both switch to the mesenchymal cell state and select for RAS-MAPK pathway mutations under the pressure of intensive combination chemotherapy (6,11,43–47). Thus, the switch to the mesenchymal cell state provides immediate resistance to anti-neuroblastoma drugs but relatively indolent cell growth, allowing for the long term outgrowth of rare subclones of cells with drug resistance due to RAS-MAPK pathway mutations, and impart aggressive proliferative and metastatic growth properties, which may allow the neuroblastoma cells to shift back to the adrenergic cell state. We have already shown that homozygous *NF1* inactivating mutations confer very aggressive growth properties in the zebrafish neuroblastoma model (48). RAS-MAPK mutations are not present in low-risk TATA/TATA tumors that arise de novo with the mesenchymal cell state, and thus these neuroblastomas remain localized and are treatable with surgery alone. The hypothesis that neuroblastoma cells harbor both RAS-MAPK pathway mutations and the adrenergic pattern of gene expression at relapse after intensive chemotherapy regimens can be tested when both whole-genome DNA sequence and genome-wide RNAseq studies are available from cohorts of patients treated with intensive chemotherapy.

Our results also revealed that a subset of the GATA/TATA neuroblastomas with lower LMO1 expression can develop in neuroblastomas with the adrenergic CRC, due in part to aberrant upregulation of LMO3, a closely related LMO family member. This is the case for the neuroblastoma tumors in our UMAP analysis (cluster 2h, Fig. 6h), which are characterized by lower *LMO1* but high *LMO3* expression levels (Fig. 6 f,h). These tumors are enriched for cases with a heterozygous *GATA/TATA* genotype, and elevated *LMO3* levels apparently cooperate with *LMO1* to drive the adrenergic CRC. The redundancy of *LMO* gene family members is analogous to the findings in the TAL1-overexpressing subtype of T-ALL, in which *LMO1* or *LMO3* can be aberrantly activated by chromosomal translocation and substitute for *LMO2* in the TAL1 CRC that drives thymocyte transformation (49–51). In this regard, it is interesting that aberrantly high *LMO3* expression levels appear to be frequent in neuroblastomas with *MYCN* gene amplification (Fig 6i), which may explain the lack of association of the rs2168101 genotype with the risk of developing *MYCN*-amplified neuroblastomas (20).

### *lmo1* -/- zebrafish do not exhibit either the mesenchymal or adrenergic CRC

An unexpected finding arising from our *in vivo* zebrafish model is that neuroblastomas arising in the *lmo1-/-* background expressed genes that were not associated with either the adrenergic or the mesenchymal CRC, but instead showed upregulation of a third set of transcription factors (Fig. 5d). This finding suggests that neuroblastoma is driven by an alternative CRC when *lmo1* is inactivated in all tissues of the host, rather than just in the neuroblastoma cells. These *lmo1-/-* tumor cells also fail to show signs of NOTCH signaling, such as upregulation of *hes2* as observed in *TATA/TATA* fish (Fig. 5c), which is potentially important because NOTCH signaling is the prototypic pathway demonstrating the important of the signaling cell as well as the receiving cell for successful signal transduction. In future studies, it will be important to conduct single cell RNAseq of unsorted cells from the neuroblastoma as they develop in fish with the different rs2168101 and *lmo1* genotypes, to test the hypothesis that the TATA/TATA genotype primarily reduces *lmo1* expression in neuronal progenitor cells, while the *lmo1* knockout affects lmo1 regulated gene expression in every tissue, including adrenal medullary stromal cells.

## Conclusions

Overall, our studies support the value of zebrafish and other *in vivo* model systems to investigate the mechanisms underlying the multi-step process of clonal progression in the initiation of cancers like neuroblastoma, and to link these underlying mechanisms to GWAS-based associations with key haplotypes that predict the risk of developing specific types of human cancers.

## Materials and Methods

### Zebrafish Lines and Maintenance

Zebrafish were derived from the AB background strain. All zebrafish were maintained under standard aquaculture conditions at the Dana-Farber Cancer Institute. All experiments were approved by the Institutional Animal Care and Use Committee (IACUC) under protocol #02-107. Zebrafish lines *Tg(dβh:EGFP)* and *Tg(dβh:MYCN)* were described previously (21) and designated *EGFP* and *MYCN* in the text, respectively.

### Genome editing of *lmo1* using TALEN and CRISPR/Cas9 technology

To generate stable lines carrying the protective T allele at rs2168101 in the first intron of *lmo1*, we designed Transcription Activator-Like Effector Nuclease (TALEN) recognition sequences that bind to the first intron of *lmo1* surrounding the *GATA* site: TALEN1 5’-TACGACTGATTTGATTTT-3’ and TALEN2 5’-TTCATTTCAAGTTCCAT-3’. A 16-bp spacer with an EcoRV sequence was located between the two binding sites. TALEN expression vectors harboring a wild-type FokI nuclease were generated as previously described(52), linearized by PmeI and used as templates for TALEN mRNA synthesis using the mMessage mMachine T3 Kit (Ambion). For the targeted integration of the protective T allele, we generated a 41-nucleotide single-stranded oligonucleotide (*TATA-ssOligo*) containing the T-allele sequence with 20 flanking nucleotides on either side of the T. The *TATA-ssOligo* also contains two additional nucleotide changes that are located 5′ prime to the T (CC instead TT, marked in bold in Fig. 2A) to not only create a new binding site for TfiI (to allow for screening for offspring with successful knock-in of the *TATA-ssOligo*) but also to prevent the TALEN1 from binding and cutting out the integrated oligonucleotide. Equal amounts of TALEN1 and 2 mRNA (100 ng/μL) together with 100pg of oligonucleotide were injected together into 1-cell-stage zebrafish embryos. To identify positive founder fish with a successful TATA knock-in, germline DNA was extracted by tail clip, and the DNA fragment surrounding the TATA site was amplified using the primers (“*TATA/TATA-EcoRV”*) listed in Table S1, followed by EcoRV digestion. Successful genome editing was detectable by loss of the EcoRV restriction site (Fig. S3A). Only fish incorporating the *TATA-ssOligo* screened positive in the TfiI assay, where TfiI digestion of an amplicon generated using the primers (“*TATA/TATA-TfiI*”) listed in Table S1 resulted in two DNA fragments: an undigested 130-bp long fragment and a digested 65-bp long fragment (Fig. S3B). Using this approach, we isolated a zebrafish line with a heterozygous T allele at rs2168101 (designated “*TATA/GATA*”, Fig. 2A). F1 embryos at 24 hpf were again screened for germ-line transmission, and the F1 embryos from the positive founder were raised to adulthood. The F2 generation was crossed with the zebrafish lines *Tg(dβh:EGFP)* and *Tg (dβh:MYCN)* and then intercrossed to obtain F4 heterozygous, homozygous mutant offspring and wild-type *(GATA/GATA*) offspring in the *dβh:MYCN;dβh:EGFP* double transgenic background.

The *lmo1* knockout zebrafish line (*lmo1*-/-) was generated by using CRISPR-Cas9 genome editing technology, targeting exon 2 (53). The following *lmo1* exon 2 CRISPR site was designed using a CRISPR Design web tool (http://crispr.mit.edu): 5′-GGAGAGGGAGATCAGATCGA-3′, CRISPR gRNA was prepared by the cloning-free, single-guide RNA synthesis method ((54)) using the Ambion T7 MEGAscript Kit (AM1334M, Ambion) and purified with the Qiagen miRNeasy Kit (217004, Qiagen). *Cas9* nuclease was purchased from New England Biolabs (M0386T). Each embryo was injected with 1 nl of solution containing *lmo1*-specific gRNA (60 ng/ul) and Cas9 (30 ng/ul) at the one-cell stage. Mosaic F0 fish with germline mutations were identified, and the stable mutant line for *lmo1+/-* was established by outcrossing to wild-type AB fish. To genotype the *lmo1+/-* line at 3 months post-injection, genomic DNA was extracted from fin clips, and DNA fragments containing the mutated site were amplified using the primers (“*lmo1-/-*“) listed in Table S1. The amplified DNA product was analyzed using the T7 endonuclease I mismatch cleavage assay to detect mutations introduced after Cas9 cleavage. One line was identified with a 32-nucleotide deletion starting at coding nucleotide 165 that caused an early stop codon at the end of the first LIM domain and a complete loss of the second LIM domain (Fig. S4). This line therefore harbors a loss-of-function allele of *lmo1* (*lmo1*+/-). The F2 zebrafish were crossed with the zebrafish lines *Tg(dβh:EGFP)* and *Tg (dβh:MYCN)*, and next intercrossed to obtain F4 *lmo1*+/+, *lmo1*+/-, and *lmo1*-/- offspring in the *dβh:MYCN;dβh:EGFP* double transgenic background.

### Neuroblastoma tumor watch

*Lmo1*+/- and *TATA/GATA* lines were crossed to *dβh:MYCN* and *dβh:EGFP* lines to obtain *lmo1*+/- and *GATA/TATA* lines in both transgenic backgrounds. Offspring were sorted for EGFP and inbred to their counterparts to obtain all possible genotypes: *GATA/GATA* (wild-type for *lmo1*), *lmo1*+/-, *lmo*-/-, *TATA/GATA*, and *TATA/TATA* in the *dβh:MYCN;dβh:EGFP* double transgenic background. Fish were anesthetized with tricaine and examined every 2 weeks beginning at 5 weeks post-fertilization (wpf) for fluorescent, EGFP-expressing cell masses. Zebrafish with EGFP-positive tumors were separated and subsequently monitored for neuroblastoma tumor progression. The cumulative frequency of neuroblastoma development was analyzed by the Kaplan-Meier method, and the log-rank test was used to determine statistical significance.

### RNA sequencing (RNA-Seq)

Total RNA was extracted from neuroblastomas arising in *GATA/GATA*, *TATA/TATA* and *lmo1*-/- in the *MYCN;EGFP* background using the QIAzol lysis reagent (Qiagen) and was cleaned using the RNeasy kit (Qiagen). Samples were treated with the TURBO DNase (TURBO DNA-free Kit; Ambion) and cleaned using the RNeasy MinElute Cleanup kit (Qiagen). Strand-specific library construction and Illumina HiSeq sequencing of paired-end 100-bp-long reads were performed at the DFCI Molecular Biology Core Facilities. All RNA-Seq data has been deposited in the GEO database (GSE No: pending). RNA-Seq data analysis was performed as previously described (55). RNA-Seq reads were aligned to the hg19 revision of the human reference genome using TopHat (55). Total mapped reads were quantified using HTSeq-count version 0.6.1. Differential expression analysis was conducted using the R Bioconductor package DESeq2 version 1.12.4, and normalized expression values for individual samples were obtained from DESeq2 using the variance-stabilizing transformation of the raw counts.

### Gene set variation (GSVA) analysis

Gene set variation analysis (GSVA) was performed in R (version 4.0.2) using the GSVA library (version 1.38.2). For each gene set of interest, scores were computed using the “kcdf=Poisson” setting with raw read counts from the Gabriella Miller Kids First neuroblastoma RNA-sequencing cohort as input. Specifically, adrenergic and mesenchymal pathway activation scores were calculated using gene lists as previously defined by Von Groningen *et al.*(6).

### Uniform Manifold Approximation and Projection (UMAP) analysis

Uniform Manifold Approximation and Projection (UMAP) analysis was performed in R (version 4.0.2) using the Seurat library (version 4.0.0). Raw FPKM data were first log base 2-transformed and scaled using default Seurat parameters. Principal components were then computed and scored using the jackstraw procedure. Ten principal components (corresponding to all components with nominal jackstraw P-value < 0.05) were then provided as input for 2-dimensional UMAP embedding. To overlay quantitative features including GSVA pathway activation scores or log base 2-transformed FPKM gene expression scores, we first computed the Z-score transform of the corresponding feature. To compute contour plots over the UMAP embedding space, we applied a Z-score-weighted Gaussian kernel to each patient datapoint that was radially symmetric with a corresponding variance equal to one half.

### rs2168101 evolution and conservation analysis

Allelic frequencies of rs2168101 across human populations were obtained from the 1000 Genomes Project Phase 3 data at the following URL: http://useast.ensembl.org/Homo_sapiens/Variation/Population?v=rs2168101. Multiple species alignment of 40 reference genomes surrounding the rs2168101 GATA site were made based on publicly available datasets on ENSEMBLE and the UCSD Genome Browser. A corresponding phylogenetic tree based on these cross-species sequences was created using a publicly available source: https://www.phylogeny.fr (56).

### Quantification and statistical analysis

Statistical analysis was performed using GraphPad Prism software version 8.0 (La Jolla, CA). Kaplan-Meier curves and log-rank tests were used to assess the rate of tumor development and differences in the cumulative frequency of neuroblastoma between fish with the following genotypes: 1) *dβh:MYCN; dβh:EGFP*, *lmo1+/+*, 2) *dβh:MYCN; dβh:EGFP*, *lmo1*+/-, 3) *dβh:MYCN; dβh:EGFP*, *lmo1*-/-, 4) *dβh:MYCN; dβh:EGFP*, *TATA/GATA*, and 5) *dβh:MYCN; dβh:EGFP*, *TATA/TATA*. Significance was determined using unpaired (two-tailed) t-tests as outlined in the figure legends. A *p*-value of less than 0.05 was considered statistically significant. For all experiments with error bars, the standard error of the mean was calculated, and the data were presented as mean ± SD. The sample size for each experiment and the replicate number of experiments were included in the figure legends.

### Data Availability

RNA-Seq data from this study is available through GEO per accession number: pending.

### Web Resources

- Geography of Genetic Variants Browser, http://popgen.uchicago.edu/ggv/

- Phylogenetic tree, https://www.phylogeny.fr

- UCSC Genome Browser, https://genome.ucsc.edu/

- depmap.com

- Gabriella Miller Kids First (GMKF) Pediatric Research Program: https://commonfund.nih.gov/kidsfirst/

## Acknowledgments

We thank members of the Look, Maris, and Young laboratories for helpful scientific discussions. We thank Cicely Jette for helpful editorial comments on the manuscript.

## Funding

This work was supported by grants from the National Cancer Institute, National Institute of Health, R35 CA210064 (A.T.L), R01 CA180692 (J.M.M/A.T.L.), R35 CA220500 (J.M.M.), R01 GM088040 (J.K.J.), K08 CA245251 (A.D.D), T32 CA009140 and R38 HL143613 (D.A.O.), and T32 HL007574-39 (N.W.L). This work was also supported by the St. Jude Children’s Research Hospital Collaborative Research Consortium on Chromatin Regulation in Pediatric Cancer (ATL, RAY, ADD, and BJA). N.W.L. is supported by the German Cancer Aid. A.D.D. and M.W.Z are Damon Runyon–Sohn Pediatric Fellows supported by the Damon Runyon Cancer Research Foundation, DRSG-24-18 (A.D.D.), DRSG-9-14 (M.W.Z.). A.D.D. and M.W.Z are recipients of Alex’s Lemonade Stand Foundation Young Investigator Awards. A.D.D. is the recipient of funding from the Rally Foundation for Childhood Cancer Research and CureSearch for Children’s Cancer Foundation. M.W.Z. is supported by the Claudia Adams Barr Program for Innovative Cancer Research. BJA is the Hope Funds for Cancer Research Grillo-Marxuach Family Fellow. A.D.D. and B.J.A. are supported by funding from the American Syrian Lebanese Associated Charities (ALSAC).

## Author Contributions

NWL, JMM and ATL conceived the study and designed the experiments. NWL, TT, HS, ADD, and MWZ performed zebrafish experiments and gene expression assays. DAO, ACW and SD analyzed human gene expression assays. DAO performed the phylogenetic tree analysis and the analysis of the human neuroblastoma gene expression data. SH and JKJ created the TALENs used in this study. BJA and RAY performed the analysis of zebrafish gene expression. NWL and ATL wrote the manuscript with input from each of the authors.

## Competing Interests

RAY is a founder and shareholder of Syros Pharmaceuticals, which is discovering and developing therapeutics directed at transcriptional pathways in cancer. BJA is a shareholder of Syros Pharmaceuticals. ATL is a founder and shareholder of Light Horse Therapeutics, which is discovering and developing small molecules to disrupt oncogenic protein complexes. JMM is a founder of Tantigen BIO and HULA Therapeutics, two companies focused on discovery and development of immunotherapies for childhood cancers. No other potential conflicts of interest are declared. J.K.J. has financial interests in Beam Therapeutics, Chroma Medicine (f/k/a YKY, Inc.), Editas Medicine, Excelsior Genomics, Pairwise Plants, Poseida Therapeutics, SeQure Dx, Inc., Transposagen Biopharmaceuticals, and Verve Therapeutics (f/k/a Endcadia). J.K.J.’s interests were reviewed and are managed by Massachusetts General Hospital and Partners HealthCare in accordance with their conflict-of-interest policies. J.K.J. is a co-inventor on various patents and patent applications that describe gene editing and epigenetic editing technologies.

## Data and materials availability

All RNA-Seq data has been deposited in the GEO database (GSE No: pending). Neuroblastoma RNA-seq available through the Gabriella Miller Kids First (GMKF) Pediatric Research Program: https://commonfund.nih.gov/kidsfirst.

## Notes

http://popgen.uchicago.edu/ggv/

https://genome.ucsc.edu/

https://depmap.org/portal/

https://commonfund.nih.gov/kidsfirst/

