## Supplemental Materials and Data for "Genetic Predisposition to Neuroblastoma Results from a Regulatory Polymorphism that Promotes the Adrenergic Cell State"

### Supplementary Materials

Figure S1

```
Zebrafish               AAA-GGGTAC-GACTGATTTGATTTTTATGATATCCACAAAAATGGAACTTGAAATGAAG
Zig-zag eel             AAAAGTTAGCAGACTGATGTGATTTTTTTGATAGCTGTAAAGACGGAAGTCTAAAT--GG
Spotted gar             AAAGGGTTACCACGTTGATTTGATTTTTCGATAGCTGTAAAAATGGAACTCGAAAT--GG
Red-bellied piranha     AAA-GGGTACCATGTTGATTTGATTTTTTGATAGCTGCAAAAATGGAACTTGAAATGGGG
Opossum                 AAAGGTTTGCAGGGCAGGCTTGATTTTTAGATAACTCTAAAGTGGAT-------CCCAAA
Coelacanth              AAAGGATTATAGTGCACATTTGATTATTAGATAGTGCTGAAATGGAT-------CCCAAA
Anole lizard            AAAGGATTACCGTGTAGATTTGATTTTTAGATAACACTAAAATGGAT-------CTCAAA
Turkey                  AAAGGATTACCATGTAGATTTGATTTTTAGATAACACTAAAATGGAT-------CCCAAA
Zebra Finch             AAAGGATTACCATGTAGATTTGATTTTTAGATAACACTAAAATGGAT-------CCCAAA
Rat                     AAAGGATTACCATGTAGATTTGATTTTTAGATAACGCTAAAATGGAT-------CCCAAA
Ryukyu mouse            AAAGGATTACCATGTAGATTTGATTTTTAGATAACGCTAAAATGGAT-------CCCAAA
Shrew mouse             AAAGGATTACCATGTAGATTTGATTTTTAGATAACGCTAAAATGGAT-------CCCAAA
Mouse                   AAAGGATTACCATGTAGATTTGATTTTTAGATAACGCTAAAATGGAT-------CCCAAA
Algerian mouse          AAAGGATTACCATGTAGATTTGATTTTTAGATAACGCTAAAATGGAT-------CCCAAA
Chinese hamster         AAAGGATTACCATGTAGATTTGATTTTTAGATAACACTAAAATGGAT-------CCCAAA
Prairie vole            AAAGGATTACCATGTAGATTTGATTTTTAGATAACGCTAAAATGGAT-------CCCAAA
Gibbon                  AAAGGATTACCATGTAGATTTGATTTTTAGATAACACTAAAATGGAT-------CCCAAA
Elephant                AAAGGATTACCATGTAGATTTGATTTTTAGATAACACTAAAATGGAT-------CCCAAA
Guinea Pig              AAAGGATTACCATGTAGATTTGATTTTTAGATAACACTAAAATGGAT-------CCCAAA
Horse                   AAAGGATTACCATGTAGATTTGATTTTTAGATAACACTAAAATGGAT-------CCCAAA
Marmoset                AAAGGATTACCATGTAGATTTGATTTTTAGATAACACTAAAATGGAT-------CCCAAA
Bonobo                  AAAGGATTACCATGTAGATTTGATTTTTAGATAACACTAAAATGGAT-------CCCAAA
Pig                     AAAGGATTACCATGTAGATTTGATTTTTAGATAACACTAAAATGGAG-------CCCAAA
Sheep                   AAAGGATTACCATGTAGATTTGATTTTTAGATAACACTAAAATGGAT-------CCCAAA
Dog                     AAAGGATTACCATGTAGATTTGATTTTTAGATAACACTAAAATGGAT-------CCCAAA
Dingo                   AAAGGATTACCATGTAGATTTGATTTTTAGATAACACTAAAATGGAT-------CCCAAA
Cat                     AAAGGATTACCATGTAGATTTGATTTTTAGATAACACTAAAATGGAT-------CCCAAA
Leopard                 AAAGGATTACCATGTAGATTTGATTTTTAGATAACACTAAAATGGAT-------CCCAAA
Rabbit                  AAAGGATTACCATGTAGATTTGATTTTTAGATAACACTAAAATGGAT-------CCCAAA
Goat                    AAAGGATTACCATGTAGATTTGATTTTTAGATAACACTAAAATGGAT-------CCCAAA
Cow                     AAAGGATTACCATGTAGATTTGATTTTTAGATAACACTAAAATGGAT-------CCCAAA
Mouse Lemur             AAAGGATTACCATGTAGATTTGATTTTTAGATAACACTAAAATGGAT-------CCCAAA
Vervet-AGM              AAAGGATTACCATGTAGATTTGATTTTTAGATAACACTAAAATGGAT-------CCCAAA
Olive baboon            AAAGGATTACCATGTAGATTTGATTTTTAGATAACACTAAAATGGAT-------CCCAAA
Chimpanzee              AAAGGATTACCATGTAGATTTGATTTTTAGATAACACTAAAATGGAT-------CCCAAA
Gorilla                 AAAGGATTACCATGTAGATTTGATTTTTAGATAACACTAAAATGGAT-------CCCAAA
Crab-eating macaque     AAAGGATTACCATGTAGATTTGATTTTTAGATAACACTAAAATGGAT-------CCCAAA
Macaque                 AAAGGATTACCATGTAGATTTGATTTTTAGATAACACTAAAATGGAT-------CCCAAA
Orangutan               AAAGGATTACCATGTAGATTTGATTTTTAGATAACACTAAAATGGAT-------CCCAAA
Human                   AAAGGATTACCATGTAGATTTGATTTTTAGATAACACTAAAATGGAT-------CCCAAA
```

**Figure S1:** Sequence of nucleotides in the first intron of *LMO1* adjacent to the G allele at the rs2168101 in 40 species including zebrafish (on top) and human (on bottom). Grey box marks the *GATA* motif.

**Figure S2**

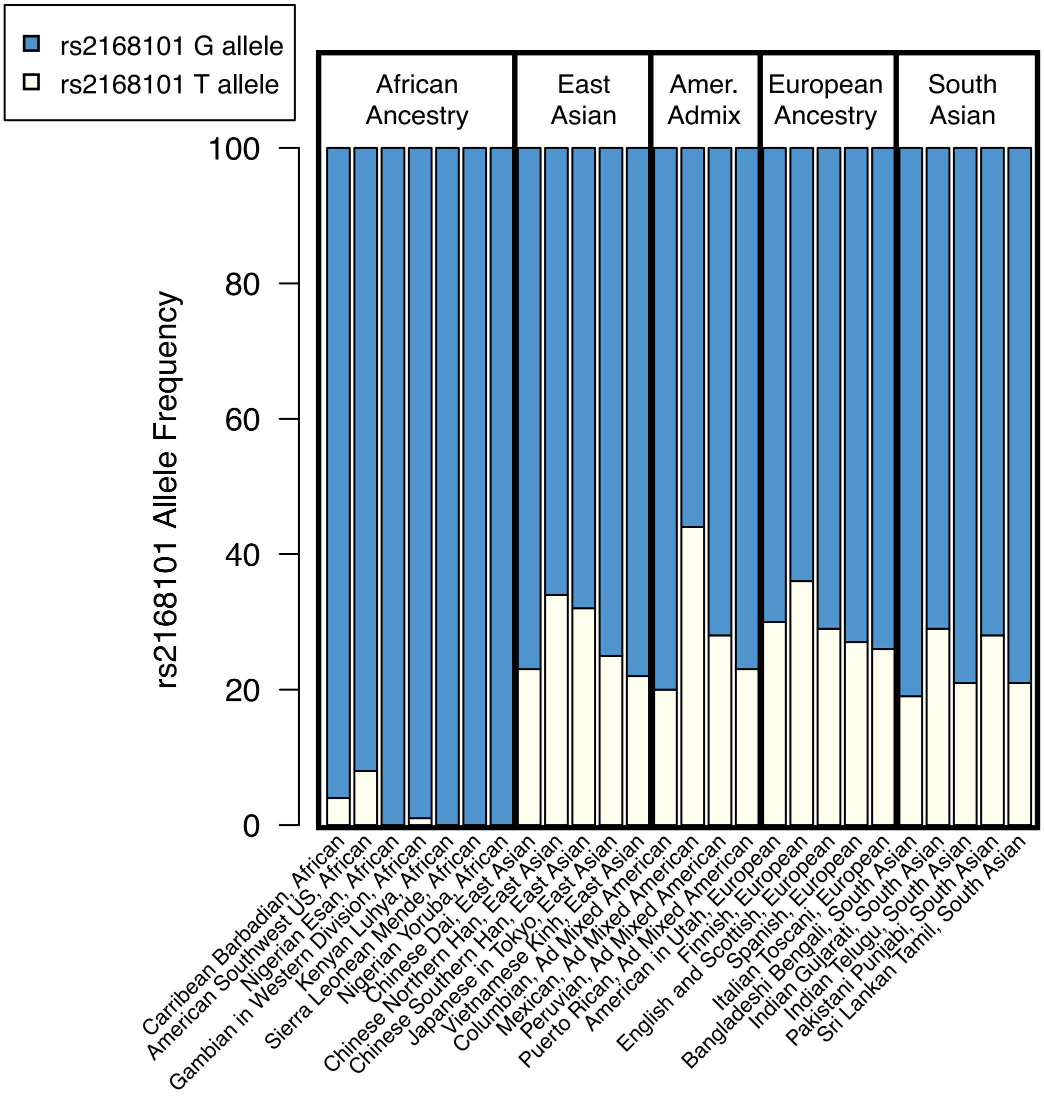


**Figure S2:** Frequency of G and T alleles at rs2168101 within geographically defined human populations genotyped as part of the 1000 Genomes Project as indicated. Frequency of the G allele is marked in blue and of the T allele in white.

**Figure S3**

**A**

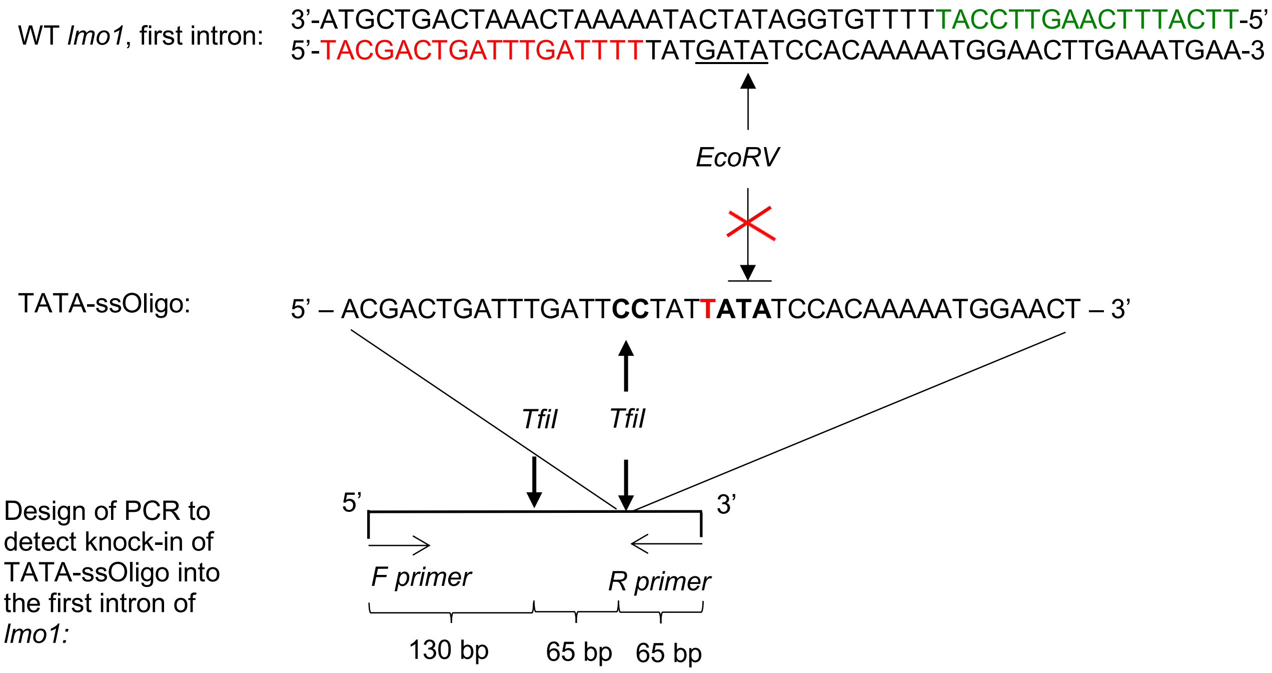


**B**

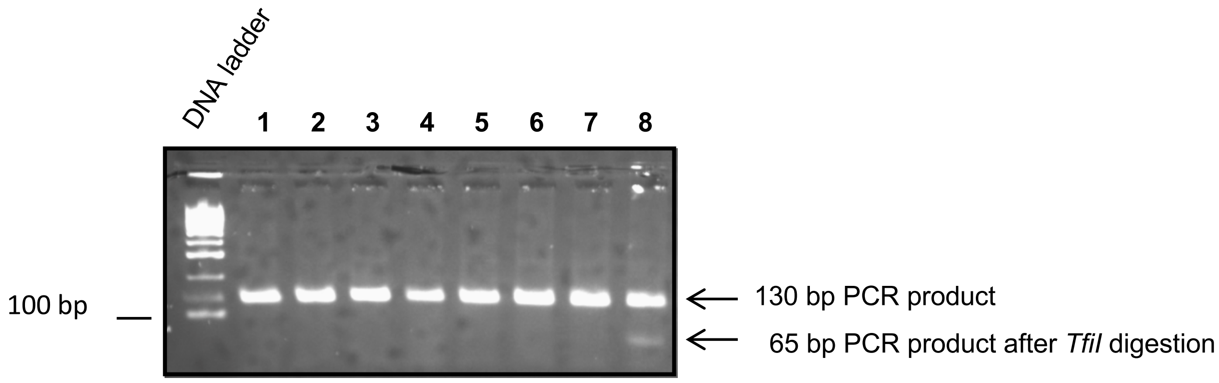


**Figure S3:** Strategy for TALEN-mediated genome editing to introduce the rs2168101 T allele in zebrafish.

(A). Wild-type (WT) sequence of the first intron of *lmo1* flanking to the G allele at rs2168101 in zebrafish is shown. The TALEN binding sites are marked in red (TALEN1) and green

(TALEN2), and the *GATA* site is underlined. The arrow points to the EcoRV restriction enzyme site that is present in the G allele but absent in the T allele. *TATA-ssOligo* is a 41-nucleotide, single-stranded DNA oligonucleotide that was co-injected with the sequence-specific TALENs into fertilized wild-type zebrafish embryos at the one-cell stage of development to facilitate knock-in of the T at rs2168101 (marked in red). The *TATA-ssOligo* also includes two other nucleotide changes ("CC", marked in bold) on the 5′ side of the T, which were included to prevent unnecessary recurrent TALEN binding (see TALEN binding sites underlined above in the wild-type *lmo1* sequence) and to create a new TfiI restriction site for screening purposes. A PCR/Tfi1 digestion assay was designed to screen zebrafish embryos for knock-in of the *TATA* ss-Oligo into the first intron of *lmo1* (below).

(B). In the PCR/Tfi1 digestion assay, only embryos with successful knock-in of the rs2168101 T allele reveal a PCR product that can be Tfi1-digested into two 65-bp products.

**Figure S4**

## A

*lmo1*

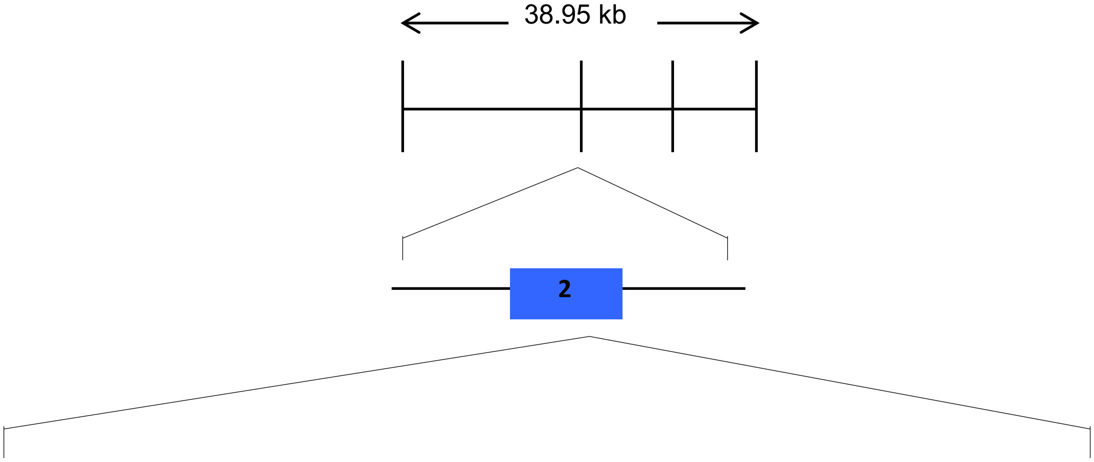


5' – GTGCCTGCTGTGACTG**CCG**CCTGGGGGAGGTGGGCTCCACCCTCTACACCAAAGCCAATCTCAT– 3'

## B

lmo1 exon 2 …TGACTG**CCG**CCTGGGGGAGGTGGGCTCCACCCTCTACACCAAAGCCAATCTCAT…
lmo1 Δ32 (-32) …TGACTG**CC** - - - - - - - - - - - - - - - - - - - - - - - - - - - - - - - AAAAGCCAATCTCAT…

## C

Lmo1
MVLDKEEGVPMLSVQPKGKQKGCAGCNRKIKDRYLLKALDKYWHEDCLKCACCDCRLGEVGSTLYTKANLIL
CRRDYLRLFGTTGNCAACSKLIPAFEMVMRARDNVYHLDCFACQLCNQRFCVGDKFFLKNNMILCQMDYEEG
QLNGSFETQVQ
Lmo1 Δ32
MVLDKEEGVPMLSVQPKGKQKGCAGCNRKIKDRYLLKALDKYWHEDCLKCACCDC***QSQSHPLSQGLPEAL***
***WYNGELCSLQX***

## D

Human LMO1 (158 a.a.)

Zebrafish Lmo1 (155 a.a.)

Zebrafish LMO1Δ32 (81 a.a.)

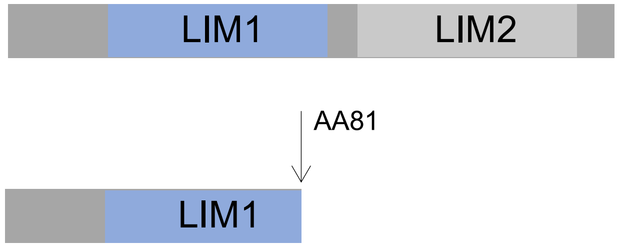

**Figure S4:** Strategy for genome editing to disrupt the coding sequence of lmo1 and generate null alleles in zebrafish.

(A). A diagram of the lmo1 gene is shown with expansion of its second exon below. The sequence recognized by the single guide RNA (sgRNA) in the coding sequence of lmo1 exon 2 is marked in green and underlined, with the Cas9 recognition site marked in red.

(B). The WT sequence of the first intron of lmo1 in zebrafish with the marked CRISPR/Cas9 site is shown in top panel. The sequence of the identified founder fish with a 32-nucleotide deletion ("lmo1 Δ32 (-32)") is shown below. The resulting knockout line is referred to as "lmo1-/-".

(C). The wild-type zebrafish Lmo1 protein sequence is shown above, with the corresponding protein sequence of the lmo1 Δ32 knockout mutant shown below with the early stop codon (red X).

(D). An illustration of the wild-type zebrafish Lmo1 protein is shown above with the LIM1 domain marked in blue and the LIM2 domain in green. The protein encoded by the lmo1 Δ32 mutant gene is shown below. The Δ32 deletion causes a frame shift and an early stop codon that truncates the first LIM domain and completely removes the second LIM domain.

**Table S1**

| zebrafish genes | Forward primer (5'-3') | Reverse primer (5'-3') |
|---|---|---|
| *lmo1-/-* | TTTCCAGACACAAGGACAAGGATCC | GCAGATATCAGAAGAAAGAGTACATTTGTCTT |
| *TATA/TATA- Tfi1* | GCTTTCAACTGTGTGCTCATTAAAAG | GATAGGATCCGGTTTCTATGTAGGC |
| *TATA/TATA-EcoRV* | GACCCTGCATACAGTAAAGTTTGCA | ATCCGCCTCTTTTCCGCCCATCCAC |
